# Abstract neural representations of category membership beyond information coding stimulus or response

**DOI:** 10.1101/2020.02.13.947341

**Authors:** Robert M. Mok, Bradley C. Love

## Abstract

For decades, researchers have debated whether mental representations are symbolic or grounded in sensory inputs and motor programs. Certainly, aspects of mental representations are grounded. However, does the brain also contain abstract concept representations that mediate between perception and action in a flexible manner not tied to the details of sensory inputs and motor programs? Such conceptual pointers would be useful when concept remain constant despite changes in appearance and associated actions. We evaluated whether human participants acquire such representations using functional magnetic resonance imaging (fMRI). Participants completed a probabilistic concept learning task in which sensory, motor, and category variables were not perfectly coupled nor entirely independent, making it possible to observe evidence for abstract representations or purely grounded representations. To assess how the learned concept structure is represented in the brain, we examined brain regions implicated in flexible cognition (e.g., prefrontal and parietal cortex) that are most likely to encode an abstract representation removed from sensory-motor details. We also examined sensory-motor regions that might encode grounded sensory-motor based representations tuned for categorization. Using a cognitive model to estimate participants’ category rule and multivariate pattern analysis of fMRI data, we found left prefrontal cortex and MT coded for category in absence of information coding for stimulus or response. Because category was based on the stimulus, finding an abstract representation of category was not inevitable. Our results suggest that certain brain areas support categorization behaviour by constructing concept representations in a format akin to a symbol that differs from stimulus-motor codes.

## Introduction

Concepts organize our experiences into representations that can be applied across domains to support higher-order cognition. How does the brain organize sensory input into an appropriate representation for categorization? Are concepts simply a combination of sensory signals and motor plans, or does the brain construct a separate concept representation, abstracted away from sensory-motor codes? Despite much research on how people organize sensory information into a format suited for categorization (e.g. Nosofsky, 1986; Kruschke, 1992; Love et al., 2004) and its neural basis (e.g. Bowman & Zeithamova, 2018; Cromer et al., 2010; Davis et al., 2012b, 2012a; Folstein et al., 2013; Freedman & Assad, 2006; Mack et al., 2013, 2016; Seger & Miller, 2010; Sigala & Logothetis, 2002; Zeithamova et al., 2019), few have explicitly examined whether category representations exist independently of sensorymotor information (Figure 1A).

**Figure 1.**
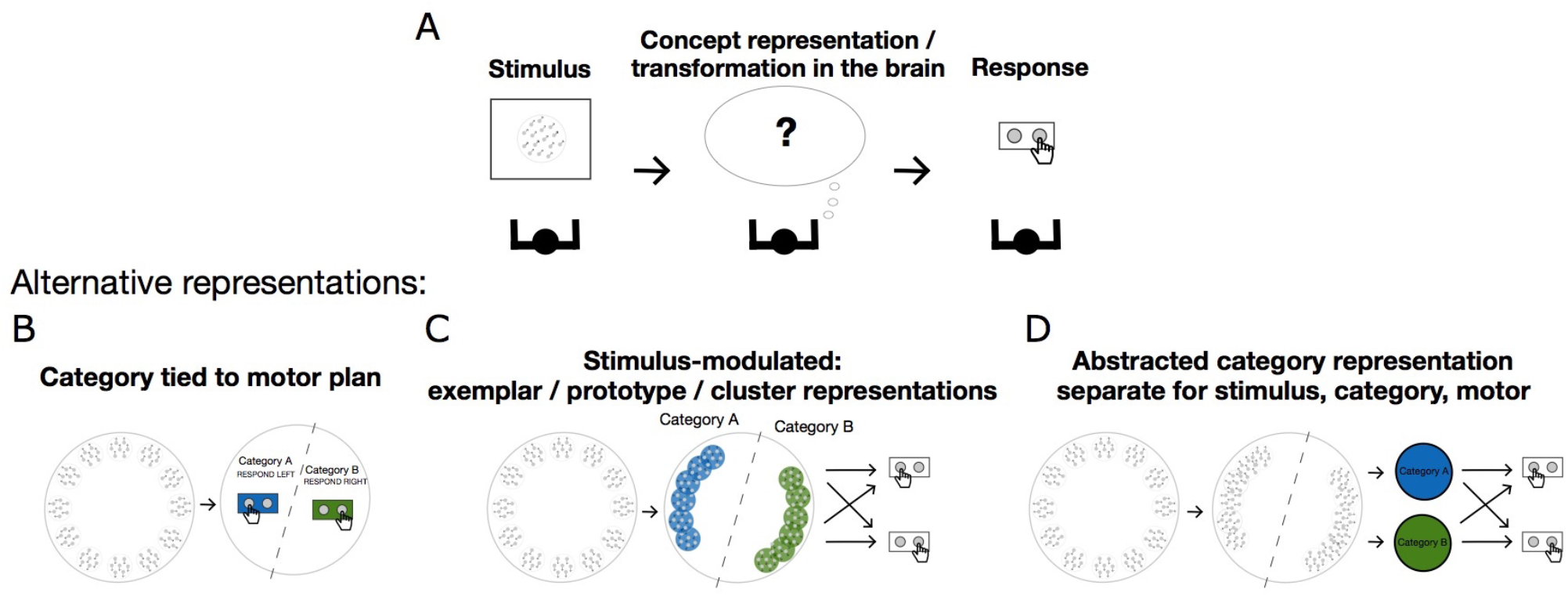
How the brain transforms stimulus into a concept representation for categorization. Stimuli are 12 motion dot patterns (100% coherent), from 0° to 330° in 30° steps. Blue and green colors denote the two categories. A) An observer must transform the percept into intermediate representations for accurate categorization behavior. B-D) Possible representations the brain might use for categorization. B) Each stimulus is associated with a motor response, where the category representation is grounded in sensory-motor codes. C) Stimulus-modulated representations as category representation. The stimulus representation is modulated by the category structure, which is turned into a motor representation for the response. D) The category-modulated stimulus representation is associated with an abstract representation of each category with a different representational format to the sensory motor codes (blue and green circles), which is then turned into a response.

Some concepts seem to be ‘grounded’ in sensory or motor experiences (Barsalou, 2008). For instance, the idea of ‘pain’ is based on experiences of pain, and the metaphorical use of the word is presumably linked to those bodily experiences. Certain aspects of concepts are more abstracted from first-hand experience and act more like symbols or pointers, which can support flexible cognition. For example, we know water can be used to clean the dishes, but when we are thirsty, we drink it. The same object can also appear entirely different in some contexts, such as a camouflaging stick insect appearing as a leaf, or when a caterpillar changes into a butterfly. In such cases where sensory information is unreliable or exhibits changes, an amodal symbol working as an abstract pointer may aid reasoning and understanding. Cognitive science and artificial intelligence researchers discuss the use of amodal symbols – abstracted away from specific input patterns – for solving complex tasks, arguing they provide a foundation to support higher cognition (Fodor, 1975; Marcus, 2001; Pylyshyn, 1984; also see Markman & Dietrich, 2000). In contrast, theories of grounded cognition suggest that all ‘abstract’ representations are grounded in, and therefore fully explained by, sensory-motor representations (Barsalou, 1999; Harnad, 1990). Indeed, sensory-motor variables and categories are often correlated in the real world and the brain may never need to represent ‘category’ in a way that can be disentangled from perception and action.

Here, we consider several competing accounts. Closely related to ‘grounded cognition’, some researchers emphasize a central role of action for cognition (Rizzolatti et al., 1987; Wolpert & Ghahramani, 2000; Wolpert & Witkowski, 2014), such that category representations could simply consist of the appropriate stimulus-motor representations and associations (Figure 1B). An alternative view holds that category-modulated stimulus representations are key for categorization, where stimulus information is transformed into a representation suited for categorization (as in cognitive models: e.g. Kruschke, 1992; Love et al., 2004). In these models, an attention mechanism gives more weight to relevant features so that within-category stimuli become closer and across-category stimuli are pushed apart in representational space (Figure 1C). Finally, the brain may recruit an additional amodal, symbol-like concept representation (Fodor, 1975; Marcus, 2001; Newell, 1980; Pylyshyn, 1984) to explicitly code for category, separate from sensory-motor representations. For instance, sensory information is processed (e.g. modulated by category structure), then transformed into an abstract category representation before turning into a response (Figure 1D). This representation resembles an amodal symbol in that it has its own representational format (e.g. orthogonal to sensory-motor codes), and act as a pointer between the relevant sensory signals (input) and motor responses (output). The advantage of such a representation is that it can play a role in solving the task and can persist across superficial changes in appearance and changes in motor commands. People’s ability to reason and generalize in an abstract fashion suggests the brain is a type of symbol processor (Marcus, 2001).

Here, we aimed to test whether the brain constructs an abstract concept representation separate from stimulus and motor signals, if the ‘category’ code consists of category-modulated stimulus representations and motor codes, or if it simply consists of stimulus-motor mappings. We designed a probabilistic concept learning task where the stimulus, category, and motor variables were not perfectly coupled nor entirely independent, to allow participants to naturally form the mental representations required to solve the task, and used multivariate pattern analysis (MVPA) on functional magnetic resonance imaging (fMRI) data to examine how these variables were encoded across the brain. For evidence supporting the **amodal account (Figure 1D)**, some brain regions should encode category information but not the stimulus or response. For the **category-modulated sensory account (Figure 1C)**, regions should encode both stimulus and category information, with no regions that encode category without stimulus information. Finally, for the **sensory-motor account (Figure 1B)**, regions should code for category, stimulus, and motor response (separately or concurrently), with no regions encoding category without sensory or motor information.

We recruited participants to an initial behavioral session where they first learned the task, and invited a subset of participants who performed relatively well to partake in an fMRI study. To assess how the learned concept structure is represented in the brain, we focused on brain regions implicated in flexible cognition, including prefrontal cortex (PFC) and parietal cortex, which are strong candidates for representing the abstract concept structure without being tied to sensory-motor variables, and sensorymotor regions which are involved in stimulus processing and may encode grounded representations such as category-modulated stimulus representations as the basis of concept knowledge. We focused on these regions to test for category representations *after* learning rather than testing regions that might be involved in learning (e.g. hippocampus, medial PFC), because participants spent a significant amount of time learning the category structure in a prior behavioral session (see Methods).

## Methods

### Participant Recruitment and Behavioral Session

We recruited participants to partake in a behavioral session to assess their ability to learning the probabilistic concept learning task. One hundred and thirty-one participants completed the behavioural session, and we invited a subset of higher performing participants that were MRI compatible to participate in the fMRI study. We set the threshold for being invited to no lower than 60% accuracy over two blocks of the task (50% is chance). Only two participants in the behavioral session performed below 60% accuracy.

### Participants (fMRI study)

39 participants took part in the fMRI study (most returned ~2-4 weeks after the behavioral session). Six participants were excluded due to lower than chance performance, misunderstanding the task, or falling asleep during the experiment. The remaining 33 participants (23 female) were aged 19-34 (mean: 24.04 ± 0.61 standard error of the mean; SEM). The study was approved by the UCL Research Ethics Committee (Reference: 1825/003).

### Stimuli and Apparatus

Stimuli consisted of coherently moving dots moving produced in PsychoPy (Peirce et al., 2019), images of faces and buildings (main task), and images of flowers and cars (practice). In each dot-motion stimulus there were 1000 dots, dots were 2 pixels in size, and moved at a velocity of ~0.8°per second. The dot-motion stimuli and images were 12°in diameter (or on longest axis). The fixation point was a black circle with 0.2°diameter. A grey circle (1°diameter) was placed in front of the dot stimulus but behind the fixation point to discourage smooth pursuit. The natural images were provided by members of Cognitive Brain Mapping Lab at RIKEN BSI. The task was programmed and run in PsychoPy in Python 2.7. The task was presented on an LCD projector (1024 x 768 resolution) which was viewed through a tilted mirror in the fMRI scanner. We monitored fixation with an eye tracker (Eyelink 1000 Plus, SR Research, Ottawa, ON, Canada), and reminded subjects to maintain fixation between runs as necessary.

### Behavioral Task

To examine how the brain constructs an appropriate mental representation for categorization, we designed a probabilistic concept learning task to be first performed in a behavioral session, then the same probabilistic categorization task in the fMRI session. Specifically, we set out to test whether any brain regions coded for an abstract category signal separate from stimulus and motor signals, if the category signal mainly consisted of category-modulated sensory signals, or if the category signal was simply a combination or co-existence of sensory-motor signals. To this end, we designed a probabilistic categorization task where the task variables (category, stimulus, and motor response) were not perfectly coupled, nor entirely independent.

On each trial, participants were presented with a set of moving dots moving coherently in one direction and were required to judge whether it belonged to one category (‘Face’) or another (‘Building’) with a corresponding left or right button press. The motion stimulus was presented for 1 second (s), followed by an inter-stimulus interval ranging from 1.8-7.4s (jittered), then the category feedback (Face or Building stimulus) for 1s. The inter-trial interval was 1.8s. Naturalistic images were used to encourage task engagement and to produce a strong stimulus signal.

The moving-dot stimuli spanned 12 directions from 0°to 330°in 30°steps, with half the motion directions assigned to one of two categories determined by a category bound. For half the participants, the category bound was placed at 15°, so that directions from 30°to 180°were in one category, and directions from 210°to 330°and 0°were in the other category. For the other half of the participants, the objective category bound was placed at 105°, so that directions from 120°to 270°were in one category, and directions from 0°to 90°and 300°to 330°were in the other category.

The corrective category feedback consisted of a face or building stimulus, which informed the participant which category the motion stimulus was most likely part of. The feedback was probabilistic such that the closer to the bound a stimulus was the more probabilistic the feedback was (see Figure 2A). In the practice sessions, participants were introduced to a deterministic version of the task prior to the probabilistic task (see *Experimental Procedure: Behavioral Session* section below).

**Figure 2.**
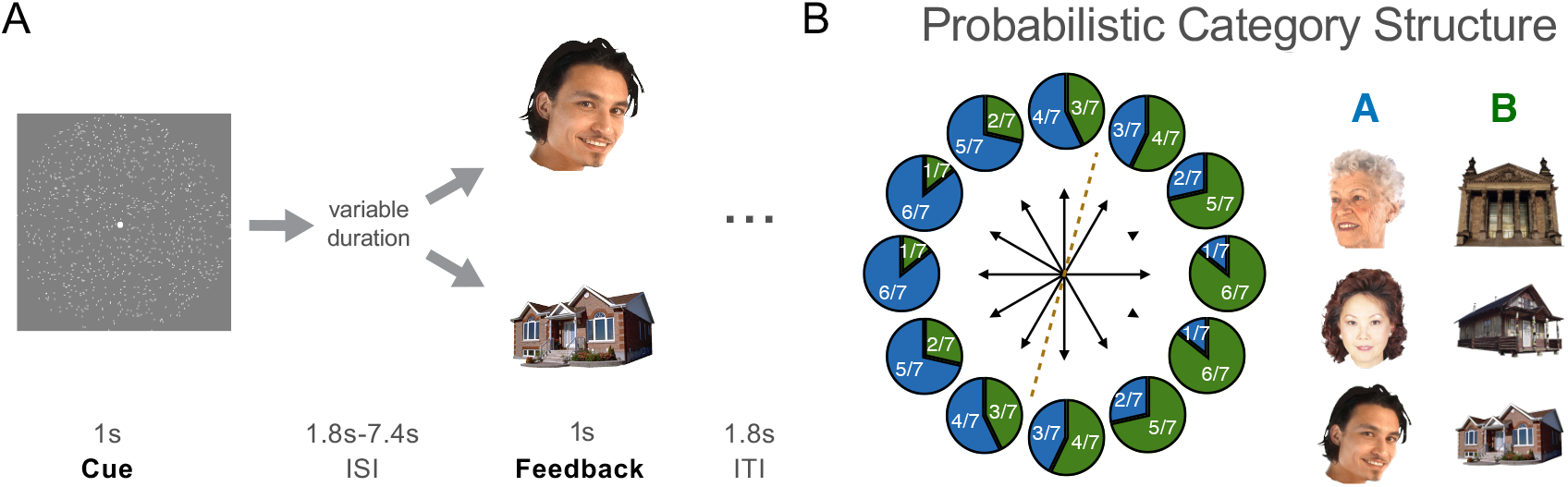
Behavioral task. A) On each trial, a dot-motion stimulus was presented and participants judged whether it was in category A or B. At the end of each trial, probabilistic category feedback (a face or building stimulus) which informed the participant which category the motion stimulus most likely belonged to. B) Probabilistic category structure. For motion stimuli to the left of the category bound (dotted line), the feedback will most likely be a face (category A), and stimuli to the right will most likely be a building (category B). For example, for the motion direction where the blue section is 4/7, the participant will see a face 4 out of 7 times, and a building 3 out of 7 times (corresponding to the 3/7 green section). The closer the motion direction to the category bound, the more probabilistic the feedback.

### Behavioral Task Rationale

Probabilistic category feedback was used in order to decouple the stimulus from the category to a certain extent. Most previous concept learning studies used deterministic feedback, such that each stimulus was always associated with the same (correct) category feedback. In terms of conditional probability, the probability of a stimulus belonging to a given category (Pr(category A | stimulus_x)) with deterministic category feedback is 1. With probabilistic category feedback, the conditional probability is less than 1, and as the stimulus-feedback association is becomes weaker (more probabilistic), it approaches 0.5 (not predictive). In this way, the stimulus and category are weakly coupled, and may lead participants to form a category representation abstracted from the more concrete experimental variables (such as stimulus and motor response). On the other hand, participants could still perform the task at greater accuracy than chance if they relied heavily on the stimulus, grounding the category in the stimulus representations.

Furthermore, the category-response association was flipped after each block (e.g. left button press for category A in the first block, right button press for category A in the second block), to discourage participants simply associating each category with a motor response across the experiment. Of course, it was still possible for participants to associate the category with a motor plan, and change this association across blocks, leading to a category representation based on motor planning.

In sum, the task required participants to learn the category that each motion-dot stimulus belonged to by its probabilistic association to an unrelated stimulus (face or building as category feedback), whereby the probabilistic feedback could lead participants to form an abstract or grounded category representation. In addition, the category-motor association was flipped across blocks. Together, the category, stimulus, and motor variables were weakly coupled, allowing us to assess whether there are brain regions that code for these variables together or independently of one another.

### Experimental Procedure: Behavioral Session (Practice and Main Experiment)

To ensure participants understood the main experimental task, they were given four practice task runs with each version gradually increasing in task complexity. In the first three runs, the task was the same as described above except that the images used for feedback were pictures of flowers and cars. In the fourth run, it was a practice run of the main task described above. Prior to each run, the experimenter explained the task to the participant.

Participants were instructed to learn which motion directions led to the appearance of Flower images and which led to the Car images. Specifically, they were told that when the moving dot stimulus appears, they should press the left (or right) button if they think a Flower will appear, or the right (or left) button if they think a Car will appear. In the first run, the category boundary was at 90° (up-down rule), and motion directions were presented in sequential order around the circle. The category (‘Flower’ or ‘Car’) feedback was deterministic such that each dot-motion stimulus was always followed by the same category stimulus feedback. For feedback, participants were presented with the image in addition to a color change in the fixation point (correct: green, incorrect: red, too slow: yellow). In the second run, the task was the same except the motion directions were presented in a random order. In the third run, participants were told that the feedback is probabilistic, meaning that the feedback resembles the weather report: it is usually it is correct, but sometimes it is not. For example, out of the 5 times you see that motion direction, you will be shown a flower stimulus as feedback 4 times, but you will be shown a car once. So the feedback is helpful on average, but sometimes it can be misleading. In the fourth run, participants were introduced to a new task to be used in the main experiment, with a new probabilistic category boundary (15° or 105°), and with face and building images as feedback.

Once participants completed the practice runs and were comfortable with the task, they proceeded to the main experimental session where they learnt the category rule from trial and error. In each block participants completed seven trials per direction condition, giving 84 trials per block. The experimenter informed participants that the category-response association flipped after each block. Participants completed three or four experimental runs.

### Experimental Procedure: fMRI Session

A subset of participants were invited to attend an fMRI session. Participants were given one practice block run as a reminder of the task, then proceeded to complete the main experiment in the scanner. Participants learned through trial-and-error. They were not informed about the location of the category boundary in either the behavioral or fMRI session, which partially explains why their performance was not at ceiling. A cognitive model fit to individuals’ behavior indicates that participants’ category boundaries differed from the optimal boundary (see below).

Participants completed three or four blocks of the probabilistic category learning task (four participants performed an extra block due to low performance on early block runs), then a motion localizer block and a face-scene localizer block (block order for localizer runs were counterbalanced across participants). After the scan session, participants completed a post scan questionnaire to assess their understanding of the task and to report their subjective category rule.

Each task block took approximately 12 minutes, and the whole scan session (main task, localizers, and structural scans) took slightly over an hour. Including preparation, practice, and post-experiment debriefing, the whole session took approximately two hours.

### Localizer Tasks

To localize the face-selective fusiform face area (FFA) (Allison et al., 1999; Kanwisher et al., 1997) and place-sensitive parahippocampal place area (PPA) (Epstein & Kanwisher, 1998) in individuals, participants completed an event-related localizer scan where they were presented with faces and buildings, and made a response when they saw an image repeat (1-back task), which was followed by feedback (the fixation point changed to green for correct and red for incorrect). On each trial, an image of a face or building was presented for 1s with inter-stimulus intervals between stimulus and feedback (green/red fixation color change) ranging from 1.8s to 7.4s (jittered), with an inter-trial interval of 1.8s. A total of 42 faces and 42 buildings were presented in a random order. Participants also completed a motion localizer run that was not used here.

### MRI data acquisition

Functional and structural MRI data were acquired on a 3T TrioTim scanner (Siemens, Erlangen, Germany) using a 32-channel head coil at the Wellcome Trust Centre for Neuroimaging at UCL. An EPI-BOLD contrast image with 40 slices was acquired in 3-mm^3^ voxel size, repetition time (TR) = 2800 ms, echo time (TE) = 30 ms, and the flip angle was set to 90°. A whole brain fieldmap with 3-mm^3^ voxel size was obtained with a first TE = 10 ms, second TE = 12.46ms, TR = 1020 ms, and the flip angle was set to 90°. A T1-weighted structural image was acquired with 1-mm^3^ voxel size, TR = 2.2 ms, TE = 2.2 ms, and the flip angle was set to 13°.

### Behavioral Model and Data Analysis

The probabilistic nature of the feedback meant that participants did not perform exactly according to the objective category rule determined by the experimenter, and inspection of behavioral performance curves suggested that most participants formed a category rule slightly different to the objective rule. To get a handle on the category rule participants formed, we applied a behavioral model to estimate each participants’ subjective category boundary from their responses.

The model contains a decision bound defined by two points, *b_1_* and *b_2_*, on a circle (0°to 359°). Category A proceeds clockwise from point *b_1_* whereas Category B proceeds clockwise from *b_2_*. Therefore, the positions of *b_1_* and *b_2_* define the deterministic category boundary between categories A and B. To illustrate, if *b_1_* = 15° and *b_2_* = 175°, stimulus directions from 15° to 175° would be in one category, and stimulus directions from 175°to 359°and from 0° to 15° would be in the other category. Note that the number of stimulus directions are not constrained to be equal across categories, as illustrated in this example (five and seven directions in each category). Despite this, there were six stimulus directions in each category for most participants.

The only source of noise in this model are the positions of *b_1_* and *b_2_* which are normally distributed as 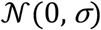. As the *σ* parameter – the standard deviation of the positions of *b_1_* and *b_2_* – increases, the position of the boundary for a given trial becomes noisier and therefore it becomes more likely that an item may be classified contrary to the position of the boundary. In practice, no matter the value of *σ*, it is always more likely that an item will be classified according to the positions of *b_1_* and *b_2_*. The standard deviation parameter provides as estimate of how uncertain participants were of the category boundary. If a participant responded perfectly consistently according to a set of bounds (deterministically), *σ* would be low, whereas if the participant was more uncertain of the bound locations and responded more probabilistically, *σ* would be higher.

The probability a stimulus *x* is an A or B is calculated according to whichever boundary *b_1_* or *b_2_* is closer. This is a numerical simplification as it is possible for the further boundary to come into play and even for boundary noise to lead to *b_1_* or *b_2_* to traverse the entire circle. However, for the values of *σ* we consider, both of these possibilities are highly unlikely. The probability that stimulus *x* is labelled according to the mean positions of *b_1_* and *b_2_* is:

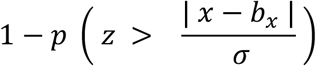

where *z* is distributed according to the standard normal distribution and *b_x_* is *b_1_* or *b_2_*, whichever is closer to *x*. Intuitively, the further the item is from the boundary position, the more likely it is to classified according to the boundary position as noise (i.e., *σ*) is unlikely to lead to sufficient boundary movement that trial. The probability an item is labeled in the alternative category (i.e., “incorrect” responses against the bound defined by the mean positions of *b_1_* and *b_2_*) is simply 1 minus the above quantity.

In other words, the probability stimulus is in a certain category is a Gaussian function of the distance to the closest bound, where the further away the stimulus is from the bounds, the more likely it is part of that category (see Figure 3A for an illustration of the model).

**Figure 3.**
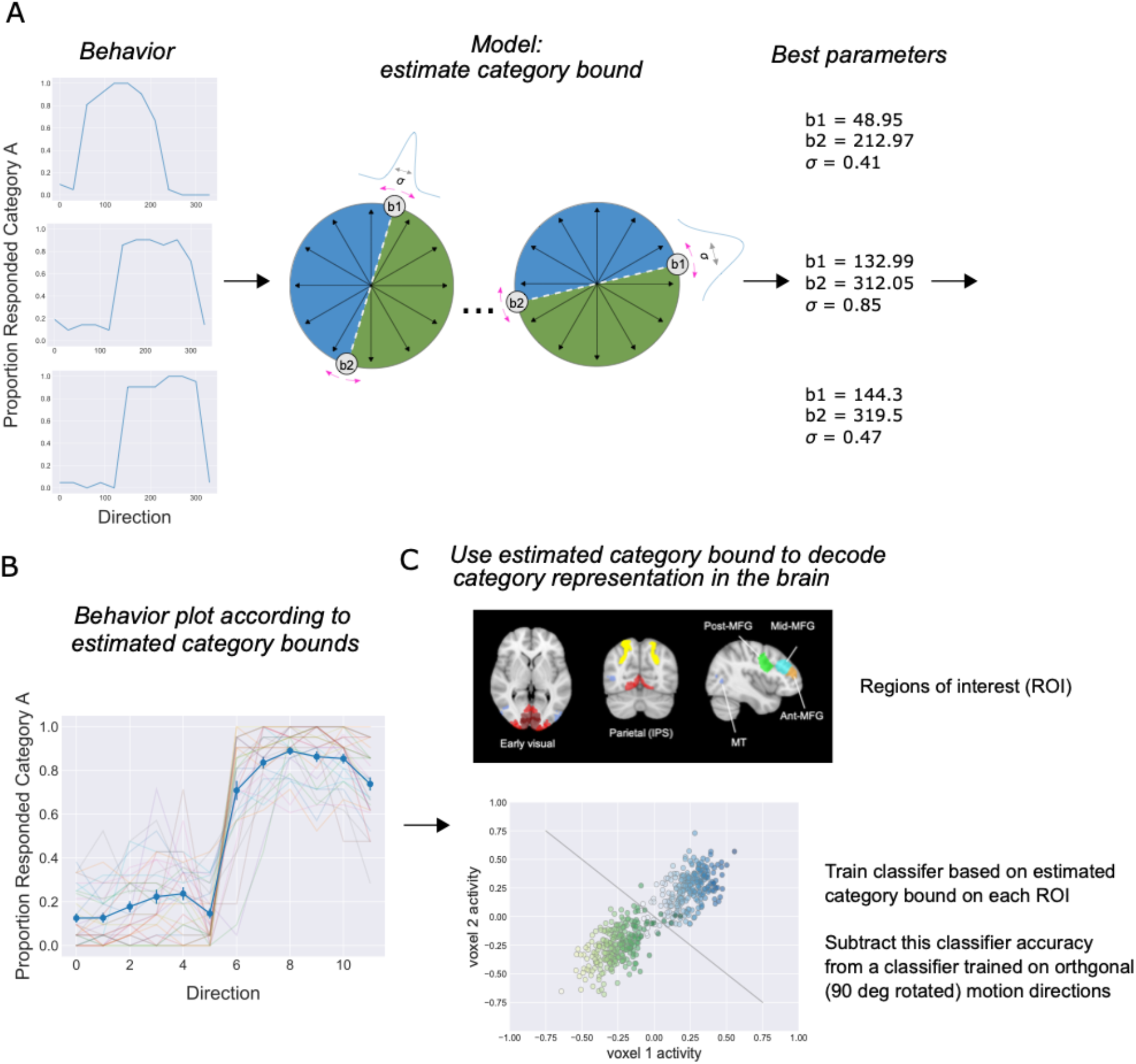
Task model, behavioral results, and model-based fMRI analysis procedure. A) The model takes individual participant behaviour as input and estimates their subjective category bound (b1 and b2) and standard deviation (σ). B) Categorization behavior. Proportion category A responses plotted as a function of motion directions ordered by individual participants’ estimated category boundary. Blue curve represents the mean, and error bars represent standard error of the mean (SEM). Translucent lines represent individual participants. C) Model-based fMRI analysis procedure illustration. Voxel activity patterns are extracted from each region of interest (ROI) for each motion direction condition (top), and a classifier was trained (support vector machine; SVM) to decode the category based on the model-based estimation of the category boundary for each participant (bottom). The data in the scatterplot were generated to illustrate example patterns of voxel activity evoked by the motion direction stimuli (two voxels shown here) belonging to each category (blue for category A, green for category B). The line is a possible support vector plane that reliably discriminates voxel patterns elicited by stimuli in category A from stimuli in category B. To test for an abstract category signal, we subtracted the classification accuracy for the category SVM by an SVM trained to discriminate orthogonal (90° rotated) directions (see Methods for details).

Maximum-likelihood estimation was used to obtain estimates for each participant (using the optimize function in SciPy). Model estimates of the subjective category bound fit participant behaviour as expected. Specifically, there was high accuracy (concordance) with respect to the estimated subjective category bound (mean proportion correct: 0.82±0.01 SEM; see Figure 3B).

Modeling and analyses was performed in Python 3.7.

### fMRI pre-processing

Results included in this manuscript come from preprocessing performed using fMRIprep 1.2.3 ((Esteban et al., 2019), RRID:SCR_016216), which is based on Nipype 1.1.6-dev ((Gorgolewski et al., 2011), RRID:SCR_002502).

#### Anatomical data preprocessing

The T1-weighted (T1w) image was corrected for intensity non-uniformity (INU) using N4BiasFieldCorrection ((B. Avants et al., 2009), ANTs 2.2.0), and used as T1w-reference throughout the workflow. The T1w-reference was then skull-stripped using antsBrainExtraction.sh (ANTs 2.2.0), using OASIS as target template. Spatial normalization to the ICBM 152 Nonlinear Asymmetrical template version 2009c ((Fonov et al., 2009), RRID:SCR_008796) was performed through nonlinear registration with antsRegistration (ANTs 2.2.0, RRID:SCR_004757, (B. B. Avants et al., 2008), using brain-extracted versions of both T1w volume and template. Brain tissue segmentation of cerebrospinal fluid (CSF), white-matter (WM) and gray-matter (GM) was performed on the brain-extracted T1w using fast (FSL 5.0.9, RRID:SCR_002823, (Y. Zhang et al., 2001)).

#### Functional data preprocessing

For each of the five or six BOLD runs found per subject (three or four task runs plus two localizer runs), the following preprocessing was performed. First, a reference volume and its skull-stripped version were generated using a custom methodology of fMRIPrep. A deformation field to correct for susceptibility distortions was estimated based on a field map that was co-registered to the BOLD reference, using a custom workflow of fMRIPrep derived from D. Greve’s epidewarp.fsl script and further improvements of HCP Pipelines (Glasser et al., 2013). Based on the estimated susceptibility distortion, an unwarped BOLD reference was calculated for a more accurate co-registration with the anatomical reference. The BOLD reference was then co-registered to the T1w reference using flirt (FSL 5.0.9, (Jenkinson & Smith, 2001)) with the boundary-based registration (Greve & Fischl, 2009) cost-function. Coregistration was configured with nine degrees of freedom to account for distortions remaining in the BOLD reference. Head-motion parameters with respect to the BOLD reference (transformation matrices, and six corresponding rotation and translation parameters) are estimated before any spatiotemporal filtering using mcflirt (FSL 5.0.9 (Jenkinson et al., 2002)). BOLD runs were slice-time corrected using 3dTshift from AFNI 20160207 (Cox & Hyde, 1997), RRID:SCR_005927). The BOLD time-series (including slice-timing correction when applied) were resampled onto their original, native space by applying a single, composite transform to correct for head-motion and susceptibility distortions. These resampled BOLD time-series will be referred to as preprocessed BOLD in original space, or just preprocessed BOLD. The BOLD timeseries were resampled to MNI152NLin2009cAsym standard space, generating a preprocessed BOLD run in MNI152NLin2009cAsym space. First, a reference volume and its skull-stripped version were generated using a custom methodology of fMRIPrep. Several confounding time-series were calculated based on the preprocessed BOLD: framewise displacement (FD), DVARS and three region-wise global signals. FD and DVARS are calculated for each functional run, both using their implementations in Nipype (following the definitions by (Power et al., 2012)). The three global signals are extracted within the CSF, the WM, and the whole-brain masks. Additionally, a set of physiological regressors were extracted to allow for component-based noise correction (CompCor, (Behzadi et al., 2007)). Principal components are estimated after high-pass filtering the preprocessed BOLD time-series (using a discrete cosine filter with 128s cut-off) for the two CompCor variants: temporal (tCompCor) and anatomical (aCompCor). Six tCompCor components are then calculated from the top 5% variable voxels within a mask covering the subcortical regions. This subcortical mask is obtained by heavily eroding the brain mask, which ensures it does not include cortical GM regions. For aCompCor, six components are calculated within the intersection of the aforementioned mask and the union of CSF and WM masks calculated in T1w space, after their projection to the native space of each functional run (using the inverse BOLD-to-T1w transformation). The head-motion estimates calculated in the correction step were also placed within the corresponding confounds file. All resamplings can be performed with a single interpolation step by composing all the pertinent transformations (i.e. head-motion transform matrices, susceptibility distortion correction when available, and co-registrations to anatomical and template spaces). Gridded (volumetric) resamplings were performed using antsApplyTransforms (ANTs), configured with Lanczos interpolation to minimize the smoothing effects of other kernels (Lanczos, 1964). Non-gridded (surface) resamplings were performed using mri_vol2surf (FreeSurfer).

Many internal operations of fMRIPrep use Nilearn 0.4.2 ((Abraham et al., 2014), RRID:SCR_001362), mostly within the functional processing workflow. For more details of the pipeline, see the section corresponding to workflows in fMRIPrep’s documentation.

### Regions of interest

To study how the brain represented category, stimulus, and response variables in the probabilistic categorization task, we focused on a set of visual, parietal, and prefrontal brain regions of interest (ROIs) hypothesized to be involved in coding these variables after learning.

We selected anatomical masks from Wang et al. (2014; https://scholar.princeton.edu/napl/resources) to examine areas involved in early visual processing, motion processing, and attention, including early visual cortex (EVC; V1, V2, and V3 merged), motion-sensitive area MT/V5 (Dubner & Zeki, 1971) and the intraparietal sulcus (IPS). We included EVC to assess stimulus-related representations including orientation and direction. The IPS is implicated in both attention (Corbetta et al., 1993; Kastner & Ungerleider, 2000; Mesulam, 1981) and category learning (Freedman & Assad, 2016; Seger & Miller, 2010). However, we did not have strong reasons to focus on specific parts of the IPS, so we merged IPS1 to IPS5 to make a large IPS ROI.

Since these masks are provided in T1 structural MRI space (1-mm^3^), when they were transformed into individual participant functional space (3-mm^3^), several masks did not cover grey matter accurately (too conservative, thereby excluding some grey matter voxels). Therefore, we applied a small amount of smoothing to the mask (with a Gaussian kernel of 0.25 mm, using fslmaths) for a more liberal inclusion of neighboring voxels, before transforming it to individual-participant space. In addition, several potential ROIs were too small to be mapped onto our functional scans. Specifically, there were several participants with zero voxels in those masks after transforming to functional space, even with smoothing. This included the motion-sensitive area MST and the superior parietal lobule (SPL1), which were not included.

Prefrontal cortex is strongly implicated representing abstract task variables (Duncan, 2001; Miller & Cohen, 2001) and task-relevant sensory signals (e.g. Erez & Duncan, 2015; Goldman-Rakic, 1995; Jackson et al., 2017; Meyers et al., 2008; Roy et al., 2010). We selected prefrontal regions implicated in cognitive control and task representations (Duncan, 2010; Fedorenko et al., 2013) http://imaging.mrc-cbu.cam.ac.uk/imaging/MDsystem) including the posterior, middle (approximately area 8), and anterior (approximately area 9) portion of the middle frontal gyrus (MFG).

Primary motor cortex was selected to examine representations related to the motor response and to test for any stimulus or category signals. Primary motor cortex masks were taken from the Harvard-Oxford atlas.

We also localized and examined brain responses in the FFA and PPA, to assess whether face and place regions, involved in processing stimuli at the feedback phase, were involved in representing the learned category (see procedure below). For example, if participants learnt that a set of motion directions belonged to category A, that was associated with face stimuli as feedback, the FFA might show information about the *learnt* category during the motion direction stimulus phase (i.e. not to the face but according to the learnt category bound). It is worth noting we are interested in assessing the information coding the *learnt* category (category A versus B), not the probabilistically presented face versus building feedback stimulus.

Apart from the FFA and PPA (where bilateral ROIs were used; see below), we included both left and right ROIs. Masks were transformed from standard MNI space to each participant’s native space using Advanced Normalization Tools (ANTs; (B. Avants et al., 2009)).

### fMRI general linear model

We used the general linear model (GLM) in FMRI Expert Analysis Tool (FEAT; Woolrich et al., 2001; FMRIB Software Library (FSL) version 6.00; https://fsl.fmrib.ox.ac.uk/fsl/) to obtain estimates of the task-evoked brain signals for each stimulus, which was used for subsequent multivariate pattern analyses (MVPA).

For the main GLM, we included one explanatory variable (EV) to model each motion stimulus trial (estimating trial-wise betas for subsequent MVPA) and an EV for each category feedback stimulus linked to each motion stimulus condition (12 EVs, not used in subsequent analyses; see trial-wise GLM examining the feedback response below). No spatial smoothing was applied. Stimulus EVs were 1s with inter-stimulus intervals between stimulus and feedback ranging from 1.8s to 7.4s (jittered), and the inter-trial interval was 1.8s. Each block run was modelled separately for leave-one-run-out cross-validation for (MVPA).

To examine motor-related brain responses, we performed an additional GLM using the same number of EVs except the EVs were time-locked to the response rather than the motion stimulus (stimulus time plus reaction time) and modelled as an event lasting 0.5s, with the assumption that the motor events were shorter than the stimulus (though this made little difference to the results). For trials without a response, the stimulus was modelled from stimulus onset as done in the main GLM, then excluded in subsequent motor-related analyses. Feedback stimuli were modelled with a single EV as above.

To localize the face-selective FFA and place-sensitive PPA, we performed an additional GLM in SPM12 (https://www.fil.ion.ucl.ac.uk/spm/software/spm12/). We applied spatial smoothing using a Gaussian kernel of full width at half maximum (FWHM) 6 mm, and included one EV for faces and one EV for building stimuli, and polynomials of degrees 0:6 to model drift in the data. Stimulus EVs were 1s with interstimulus intervals between stimulus and feedback (green/red fixation point color change) ranging from 1.8s to 7.4s (jittered), with an inter-trial interval of 1.8s. We included three contrasts Faces > Buildings, Buildings > Faces, and overall Visual Activation (Faces & Buildings). To define individual participant ROIs, we used minimum statistic conjunctions with visual activations. To localize the FFA, the conjunction was: (Face > Building) & Visual. For the PPA, the conjunction was: (Building > Face) & Visual. The rationale behind this conjunction is that functional ROIs should not only simply be selective but also visually responsive (all voxels that were de-activated by visual stimulation were not included). The conjunction was thresholded liberally at p<0.01 uncorrected. The peaks for each functional ROI was detected visually in the SPM results viewer, and we extracted the top 100 contiguous voxels around that peak.

There were four participants for which we could not find clear peaks and clusters for the left FFA, five participants for the right FFA, seven participants for the left PPA, and six participants for the right PPA. Since we were unable to reliably localize these areas for all participants in both hemispheres, we used unilateral ROIs for participants with unilateral FFA/PPA ROIs, and excluded participants for that ROI if they did not have either a left or right FFA or PPA ROI. The difficultly in localizing these areas for a subset of participants might have been due to our relatively short (two runs) event-related localizer design. In summary, when testing the FFA, we excluded two participants (no left or right FFA), and when testing the PPA, we excluded four participants (no left or right PPA).

We also performed a motion localizer, but likely due to the short localizer and the event-related design, it was not possible to reliably localize participant-specific motion-sensitive regions.

To examine information during the category feedback, performed an additional GLM modelling the same events as the main GLM (locked to motion stimuli and feedback stimuli), except that one EV was used to model each feedback trial (estimating trialwise betas for subsequent MVPA), and one EV for each motion stimulus condition (12 EVs). This additional GLM was used to estimate the trial-wise feedback mainly for practical reasons. If modelling all cue and feedback trials, it becomes a substantially larger model for FSL. By modelling the cue period trials in a separate GLM to the feedback trials, we were able to reduce the number of EVs per model (96 rather than 168).

### Multivariate pattern analysis (MVPA)

To examine brain representations of category, stimulus, and motor response, we used MVPA across our selected ROIs. Specifically, we trained linear support vector machines (SVMs; c.f. Kamitani & Tong, 2005) to assess which brain regions contained information about the category (‘Face’ or ‘Building’), stimulus (direction, orientation, and 12-way classifier), and motor response (left or right button press).

Decoding analyses were performed using linear support vector classifiers (SVC; C = 0.1) using Scikit-learn Python package (Pedregosa et al., 2011) with a leave-one-runout cross-validation procedure.

To test for abstract category coding, we first trained a classifier to discriminate between motion directions belonging to the two categories for each participant’s subjective category bound. To ensure that this was a pure category signal unrelated to stimulus differences (e.g. simply decoding opposite motion directions), we trained a classifier based on the participant’s subjective category bound, rotated 90°. For a strict test of an abstract category signal, we subtracted the classification accuracy of the first classifier from accuracy of the second classifier. The reasoning behind this is that if a brain region contains information about the stimulus direction but no information about category, it is still possible to obtain significant classification accuracy for the category classifier (category A versus category B). However, if the brain region primarily encoded stimulus information, there should be as much information for the orthogonal directions in the voxel activity patterns within an ROI (assuming sensory biases across voxels are equal). Therefore, if a brain region carries information about the category and sensory content, there would be greater classification accuracy when decoding across directions across the category boundary (with category and sensory information) than classification accuracy for the directions across the rotated boundary (sensory but no category information). The subtraction allows us to test whether the brain regions carries abstract category information over and above the sensory information. If there is only sensory information and no category information, the subtracted classification accuracies should be centred around zero. Negative values would suggest more information across directions across the boundary orthogonal to the category bound. This would most likely reflect unequal perceptual biases across voxels in that brain region, where that brain region contained more information about the motion directions across the rotated boundary compared to those across the category boundary (by chance, i.e. stimulus-based, unrelated to the learned category). Different participants were randomly assigned to different objective category boundaries, which makes a systematic bias unlikely. Previous studies have tested whether a brain region contains information that can discriminate members of different categories. However, these studies could not rule out the contributions of the stimulus features to the category decoder. In our study, stimulus feature differences are matched when comparing stimuli across the category boundary versus stimuli across the orthogonal boundary. This subtraction method ensures that the sensory signal is not the main contributor to any category code found.

To ensure this category signal was not related to motor preparation or the response, we also subtracted the former category classifier accuracy from a motor response classifier accuracy (discriminating between left versus right button presses).

For stimulus direction coding, we trained classifiers to discriminate between all six pairs of opposite motion directions (0°versus 180°, 30°versus 210°, etc.) and averaged across the classification accuracies.

To examine motor response coding, we trained classifiers to discriminate between left and right button presses on the GLM where we locked the EVs to the motor responses (reaction times).

As a control analysis, we tested whether a classifier trained on the objective category structure (i.e. defined by the experimenter) produced similar results to the subjective category analysis. The procedure was the same as the abstract category classifier above, except that the directions in each category were determined by the experimenter. In another set of control analyses, we assessed if there was any information about stimulus. We tested for orientation coding by training a classifier on all 12 pairs of orthogonal orientations irrespective of the motion direction (0°versus 90°and 0°versus 270°, 30°versus 120°, 30°versus 300°), and averaged across the classification accuracies. Finally, for a more general measure of stimulus coding, we trained a 12-way classifier to assess stimulus coding for each motion direction.

We used one-sample t-tests (one-tailed) against chance-level performance of the classifier (using SciPy, (Virtanen et al., 2020)). Multiple comparisons across ROIs were corrected by controlling the expected false-discovery rate (FDR) at 0.05 (Perktold & Seabold, 2010). For decoding category, we corrected across 12 ROIs (all apart from bilateral EVC and bilateral motor cortex), and for the direction and 12-way classifier, we corrected across 14 ROIs (excluding bilateral motor cortex). Bonferroni correction was used for tests with two ROIs (correcting for visual and motor hemispheres for orientation and motor decoding, respectively). For others, we report the uncorrected p-values since none survived even without correction. MVPA and statistical analyses were performed in Python 3.7.

### Brain-behavior correlations

To assess whether the brain’s representation of the abstract category signal contributed to categorization performance, we performed robust regression (Perktold & Seabold, 2010) to assess the relationship between categorization performance (concordance to the estimated subjective category structure) with classifier accuracy for the category for the ROIs with greater than chance classification accuracy for category.

Matplotlib (Hunter, 2007) and Seaborn (Waskom, 2020) was used for plotting and creating figures in this manuscript.

### Data and Code Availability Statement

The code for the behavioral model and data analysis will be publicly available at https://github.com/robmok/memsampCode. The behavioral and fMRI data will be made publicly available at to https://openneuro.org/

## Results

To assess how learned concept structure is represented in the brain, 33 participants learned a probabilistic concept structure in an initial behavioral session, and returned on a separate day to perform the probabilistic categorization task (the same task in the behavioral session) whilst they underwent an fMRI scan.

We used a model to estimate individual participants’ subjective category bound (see Methods and Figure 3A-B). Briefly, the model assumes that participants form a mental decision boundary in the (circular) stimulus space to separate the categories, and there is some uncertainty of the placement of this bound. Formally, the model has three parameters: the first two determines bound placement (*b_1_* and *b_2_*), and the third is a standard deviation parameter (*σ*) that models the (normally distributed) noise in this bound. *σ* provides an estimate of how certain (lower *σ*) participants are of their boundary placement. The model-estimated category bounds corresponded to participant’s categorization behavior. To compute a measure of behavioral accuracy, we the computed proportion of categorization responses consistent with individual participants’ estimated category bound. There was a strong correlation between the standard deviation parameter of the model *σ* and behavioral accuracy (r = −0.90, p = 9.00e-13), suggesting the standard deviation parameter characterizes an aspect of the categorization behavior well.

To evaluate the three main accounts of how the brain organizes information for categorization, we performed MVPA across visual, parietal, and prefrontal regions of interest (ROIs) hypothesized to be involved in representing the learned concept structure (Figure 3). Specifically, we trained linear SVMs to assess which brain regions contained information about category (A or B), stimulus (directions), and response (left or right). For a strict test of an abstract category signal unrelated to stimulus features, we trained a classifier to discriminate between motion directions in category A versus directions in category B, and subtracted this from a control classifier trained to discriminate between directions in category A rotated 90°versus directions in category B rotated 90°. This ensured that the classifier was not simply picking up information discriminating opposite stimulus directions (see Methods for details).

Our findings most strongly align with the hypothesis that the brain constructs an amodal symbol for representing category, independent of sensory-motor variables. Specifically, we found an abstract category signal over and above stimulus information in the middle portion of the left middle frontal gyrus (mMFG: p=0.0025, q(FDR)=0.029) and left motion-sensitive area MT (p=0.0086, q(FDR)=0.048; Figure 4A; 5A-B). This is particularly striking since the category is based on the stimulus direction, and there was no hint of a direction signal in these regions (p’s>0.41; Figure 4B, 5A-B).

**Figure 4.**
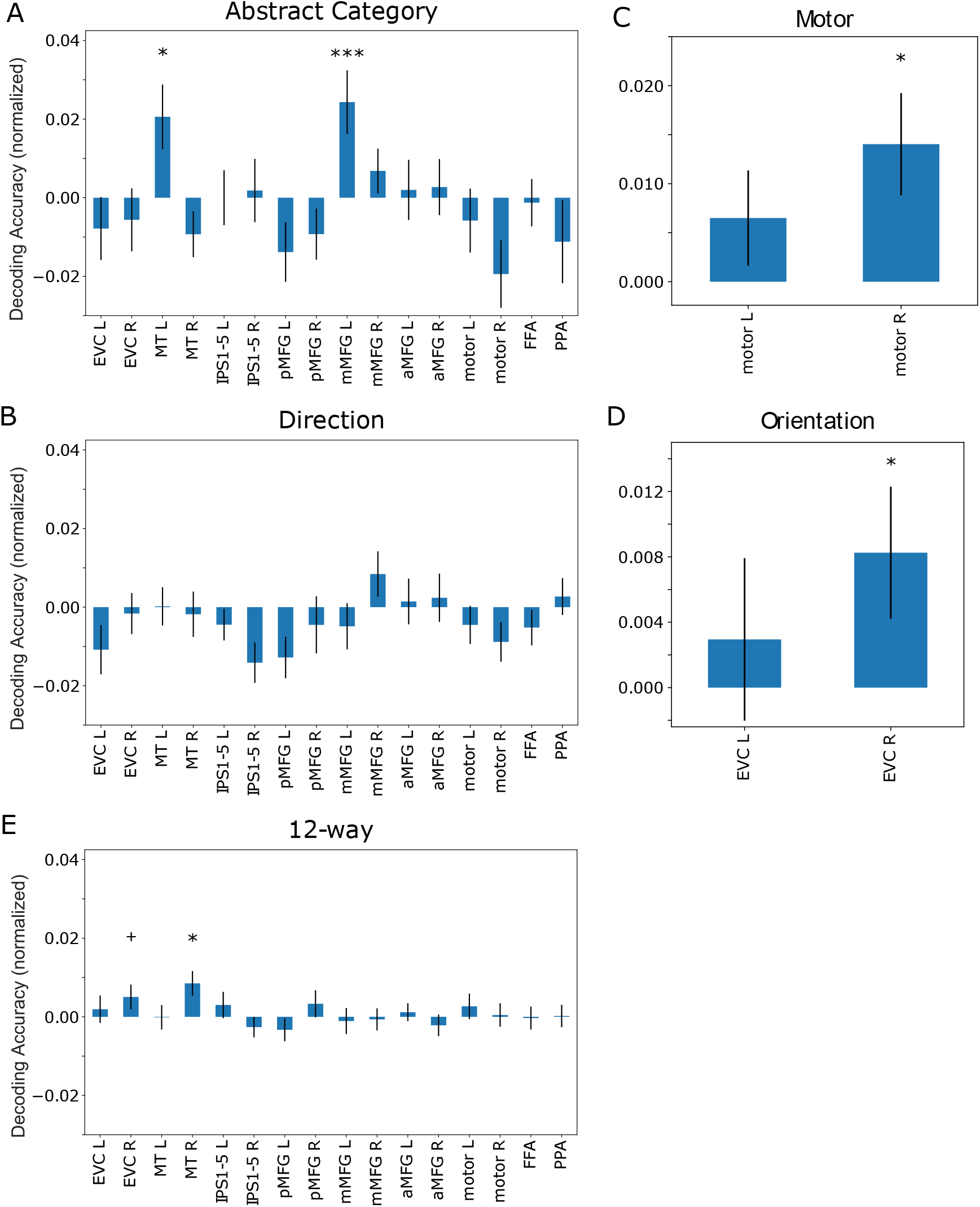
fMRI multivariate pattern analysis results. A) Abstract category coding in left mMFG cortex and left MT. Abstract category coding over and above sensory coding was computed by the category classifier accuracy minus the classifier accuracy trained on orthogonal (90 ° rotated) directions. B) No strong effects showing stimulus motion direction information. C) Right motor cortex showed significant information coding response. D) Right early visual cortex showed significant information coding orientation. E) Right MT contained sensory information as shown by the 12-way stimulus classifier, with right early visual cortex showing a similar trend. Normalized decoding accuracy measures are normalized by subtracting chance values (direction=1/2; 12-way=1/12; motor=1/2; orientation=1/2), apart from abstract category which subtracts from a control classifier (chance=0). *** p=0.0025; *p<0.05; + p<0.06.

**Figure 5.**
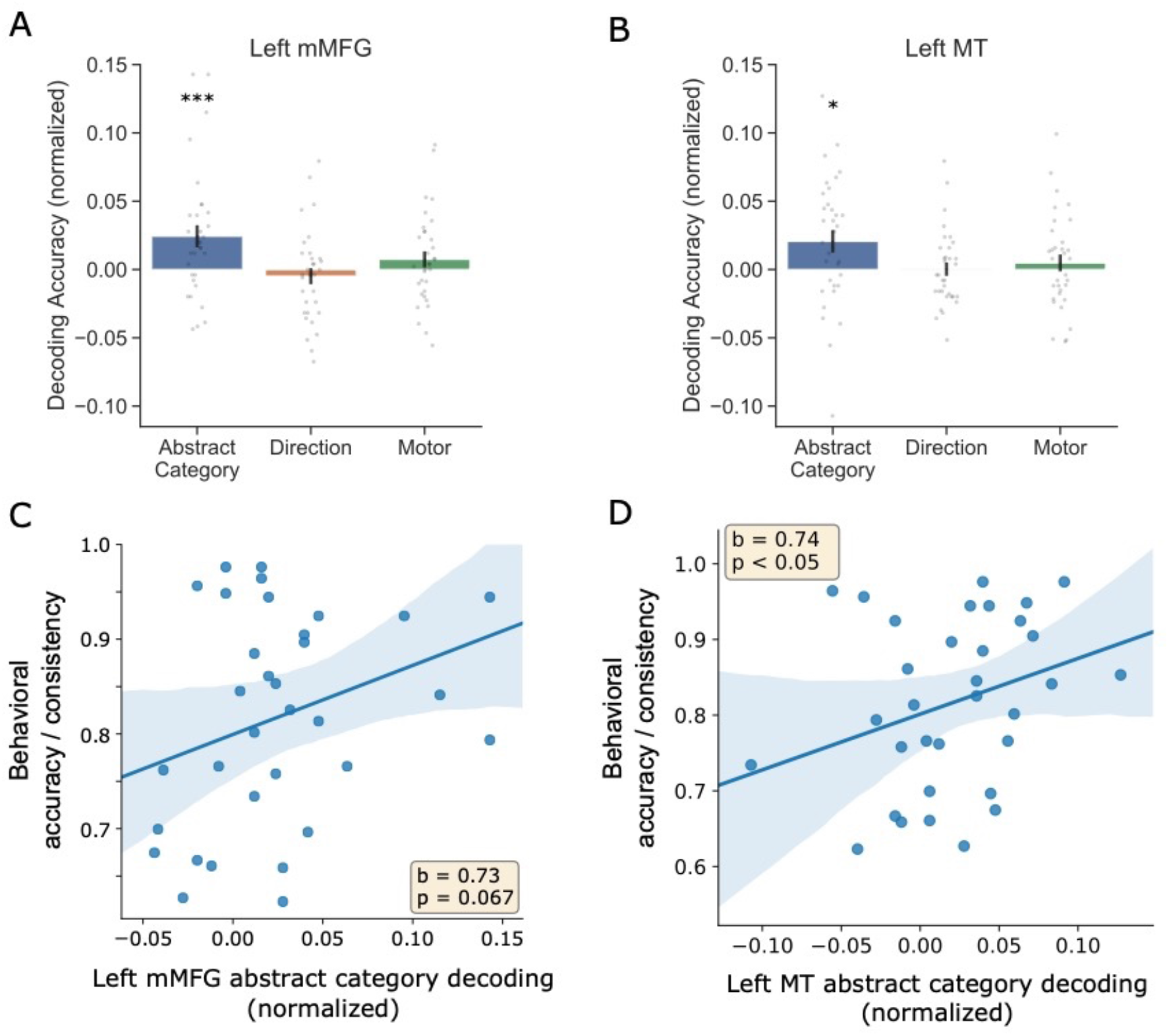
Abstract category coding and correlations with categorization behavior. A-B) Univariate scatterplots showing significant abstract category coding in left mMFG (A) and left MT (B), with no evidence of stimulus and motor coding. Grey dots are individual participants. C-D) The strength of abstract category coding in MT (D) was correlated with categorization accuracy, i.e. consistent responses with subjective category bound. There was a trend in the same direction in left mMFG (C). In D) and E), beta coefficients are from a robust regression analysis, and the shaded area represents 95% confidence intervals for the slope. *** p=0.0025, * p<0.05. Error bars represent SEM. Normalized decoding accuracy measures are normalized by subtracting chance values, apart from abstract category which subtracts from a control classifier (chance=0).

Consistent with the idea that abstract category representations can aid performance, we found that the strength of category decoding was positively correlated with categorization accuracy (responses consistent with the model-estimated category bound) in left MT (robust regression; beta=0.74, p<0.05; Figure 5D) with a similar trend for left mMFG (beta=0.73, p=0.067; Figure 5C). We also confirmed that the category signal was stronger than the motor code in left mMFG by subtracting the classifier trained to discriminate motion directions across categories from the motor classifier (p=0.015).

As expected, we found information coding motor response in motor cortex (right: p=0.006; Bonferonni-corrected for hemisphere p=0.011; left p=0.095, Bonferonni-corrected p=0.19; Figure 4C), but no information about category or direction (p’s>0.42).

Notably, abstract category coding was only present for the participant-specific subjective category structure (‘objective’ category bound classifiers across all ROIs: p’s>0.06). Furthermore, we found no evidence of category coding in the fusiform face area or in the parahippocampal place area (p’s>0.31).

Although we did not find category and stimulus representations intertwined, this was not because stimulus representations were not decodable in our data. We trained a classifier on orientation in the early visual cortex and found activity coding orientation (Figure 4D, p<0.05, Bonferroni-corrected for hemisphere). We also trained a 12-way classifier in order to assess if there was any information about the stimulus that would not be found simply by examining orientation or direction responses, and found that right MT encoded information the stimulus (p=0.005, q(FDR)=0.03), and a trend for right EVC (p=0.06, q(FDR)=0.18; Figure 4E). Notably, there was no evidence for this in the left mMFG or left MT, which encoded abstract category (p’s>0.74).

## Discussion

We examined the neural representations underlying categorization and found that the brain constructs an abstract category signal with a different representational format to sensory and motor codes. Specifically, left PFC and MT encoded category in absence of stimulus information, despite category structure being based on those stimulus features. Furthermore, the strength of this representation was correlated with categorization performance based on participants’ subjective category bound estimated by our model.

Although some representations may be grounded in bodily sensations, for tasks that require flexibility and representations to support abstract operations, an amodal symbol of a different representational format to that of sensory-motor representations may prove useful (Fodor, 1975; Marcus, 2001; Newell, 1980; Pylyshyn, 1984). Indeed, a category representation tied to a motor plan or stimulus feature would facilitate stimulus-motor representations effectively in specific circumstances, but become unusable given slight changes in context. In this study, it was possible to solve the task in multiple ways, such as a combination of the sensory-motor variables, using a category-modulated sensory representation, or additionally recruiting an amodal representation (Figure 1B-D). Despite this, we found the brain produces an additional abstract representation to support categorization (Figure 1D). Specifically, by applying a decoding approach to test for a category signal over and above a sensory code we showed that the left mMFG and MT encoded a category signal abstracted from the sensory information. Furthermore, these areas did not carry any information about stimulus or response (evidenced by the motor, direction, orientation, and 12-way stimulus classifier), and the category code was significantly stronger than the motor code. In contrast, we did not find any regions that encoded both the category and stimulus, as predicted by the account where category information is grounded in category-modulated stimulus representations (Figure 1C). We also did not find any regions that encoded both the category and the motor response, as predicted by the account where category is grounded in stimulus-motor associations (Figure 1B). Therefore, our findings suggest that brain constructs an amodal symbol for representing category, independent of sensory-motor variables. It is worth noting that our results do not suggest that the brain does not use sensory information, nor that there are no grounded neural representations, rather that the brain constructs an *additional* category representation abstracted from the sensory-motor information for categorization.

In addition to the left PFC, we found that left MT encoded a category signal in the absence of sensory information, whereas right MT was only driven by sensory information. One possible explanation is that the category signal originated from left PFC, which was sent back to modulate left MT. This may have resulted in competition between the category and sensory signals, and the task-relevant category signal won out over the bottom-up sensory signal. Since there was no category signal in right PFC, right MT was not affected by the task and coded the bottom-up stimulus signal. Alternatively, left MT may simply be more affected by top-down modulation from PFC. For instance, task-relevant attentional modulation in left PPA (when attending to scenes versus faces) seems to be stronger and more reliable than the right PPA (Chadick et al., 2014; Gazzaley et al., 2005). Unfortunately, most fMRI studies of perceptual or category learning using motion-dot stimuli did not examine left and right MT hemispheres separately, and did not report differential effects of category and stimulus across hemispheres. Future studies or meta-analytic studies could examine whether or not left or right MT is more strongly modulated by task demands, or if the lateralized modulation of sensory cortices depends on the relative lateralized recruitment of control regions such as the PFC.

Previous studies have found strong stimulus coding and category-related modulation stimulus representations during concept or perceptual learning (Braunlich & Love, 2019; Ester et al., 2020; Freedman & Assad, 2006; Kourtzi et al., 2005; Kuai et al., 2013; Mack et al., 2013; J. Zhang & Kourtzi, 2010). For example, concept learning studies that used object stimuli have shown strong modulation of sensory signals in the lateral occipital cortex after learning (Braunlich & Love, 2019; Kuai et al., 2013; Mack et al., 2013).

One major difference between prior work and the current study is the probabilistic relationship between stimulus and feedback. In the world outside the laboratory, the relationship between stimulus and feedback is not always deterministic and people must make decisions and learn in the presence of this uncertainty. For example, after viewing dark clouds and the weather forecast, a person with picnic plans is faced with the decision of whether to continue. After deciding, they update their knowledge based on whether it rained, which is a probabilistic function of what was known at the time of decision.

Another key difference between our study with many studies of concept learning is that the response mapping was switched after each block so that we could observe possible differences between category representations and stimulus-response mappings. Some researchers suggest that changing the response mapping should disrupt procedural learning processes involved in concept learning (Maddox & Ashby, 2004), which is one reason why response mappings are often held constant within a participant.

These two differences with previous studies made it possible for us to observe a strong category signal that was not strictly modulated by stimulus representations, nor motor response. This category signal was of a different format than information related to stimulus or response. We need not have observed this finding. It would have been possible for the brain to solve this task using a stimulus-modulated category representation (i.e., stimulus and category represented in related formats) in which the response mapping varied across blocks. Instead, it appears that an intermediate category signal was used by participants. Although our design did not necessitate our main finding, it is possible that the relatively loose coupling between stimulus, category, and response encouraged forming a category representation of a different format than either stimulus or response. Many real-world categories may place related demands on learners. For example, relational categories, such as thief, are not closely tied to sensory representations (Jones & Love, 2007).

It may be argued that, since participants had to flip the category-motor mapping across blocks, we encouraged participants to use the stimulus or the category information and discouraged a motor-based strategy. However, it was still possible to ground category information into motor representations by associating sensory representations to the motor plan *within* a block, and reprogram the association across blocks. Since there were relatively few blocks, this would have been possible, and a viable strategy. If the brain primarily relies on the sensory-motor association for the category representation and behavior, we would find representation of motor plans without an abstract category code (i.e. not tied to motor plans), which is not in line with our results. Furthermore, we showed evidence for an amodal category signal using a decoding approach that tested for a category signal over and above stimulus coding, and also showed that the category signal was stronger than the motor code. To test the hypothesis that our design might have discouraged participants to use motor-based representations, future studies could compare groups that had to switch the motor responses versus those that did not, and whether the latter group would form a grounded, motor-based neural representation for categorization in absence of abstract category representations.

Some of our analyses yielded negative classification accuracy values, including in the abstract category and the direction classifier accuracies (Figure 4A-B). As noted in the Methods, the purpose of subtracting the category classifier from the classifier trained on the orthogonal bound was to find category signal over and above any sensory information contained in the voxels. Negative values would simply reflect no category information, in addition to more information across directions across the boundary orthogonal to the category bound (i.e. sensory biases in the voxels unrelated to the task). For the direction classifier, there were some regions that showed negative classification accuracy values. In this analysis, the theoretically lowest possible value is zero, and values around zero would reflect the absence of any direction information. We were unable to find anything systematic that contributed to the negative values, and suggest that these effects were most likely attributable to reasons unrelated to the task, such as some non-stationarity across blocks.

There are similarities between our probabilistic conceptual learning paradigm and tasks learning the transition probability structure of object-to-object sequences. In those tasks, the probability that an object A is most likely followed by an object B (e.g. with probability of 0.75), but could also be followed by another object C (probability of 0.25) – i.e., participants learn the statistical dependencies between objects, like how our participants learn the probabilistic dependencies between stimuli and categories. Interestingly, one study by Schapiro et al. (Schapiro et al., 2013) showed participants learnt and accurately represent object-object associations with a structured community structure in several brain regions including the left prefrontal cortex. Specifically, pattern similarity analysis showed that left prefrontal cortex, anterior temporal lobe, and superior temporal gyrus encoded the statistical, relational structure across the objects. Other studies found that the regions in the medial temporal lobe including the hippocampus and entorhinal cortex are involved in the learning and retrieval of associations and prediction of object-object transitions (e.g. Garvert et al., 2017; Schapiro et al., 2016). Since our current study focused on category representations after learning, it would be interesting for future studies to test whether medial temporal lobe structures are involved in learning probabilistic conceptual structures in a similar way.

There are several open questions to be explored in the future. Our study was not optimized to study to role of the hippocampus in. category learning, as we examined category representations *after* learning. Future studies could examine the neural representations involved in probabilistic concept learning early in learning and compare them to representations during categorization after learning is complete, to explore whether how the neural representations change as concept information is consolidated into long-term memory.

Future work could also assess the causal involvement of these abstract category representations in mMFG and MT. One idea would be to use transcranial magnetic stimulation (TMS) to disrupt the left mMFG and left MT to assess whether these areas act causally support categorization behavior. It would be interesting to test whether mMFG or MT plays a more important role, by observing the fMRI signal after disruption. It could be that the mMFG is the origin of the category signal, but it is its influence on MT that leads to effective categorization behavior (e.g. TMS to mMFG leads to disruption of the MT category representation but not vice versa, where stimulation at both sites disrupt behavior).

What is the use of an abstract, symbol-like concept representation? In real-world scenarios, there are often no explicit rules and reliable feedback is rare. Building an abstract representation that can be mapped onto different contexts can be useful in real-world tasks, where the meaning of a situation can remain constant whilst the contextually appropriate stimulus or response changes. As we find here, the brain constructs an amodal, abstract representation with a different representational format separate from sensory-motor codes, well-suited for flexible cognition in a complex world.

## Acknowledgements

We thank Johan Carlin for his help on experimental design and data collection, and Amna Ali for her help on data collection. We thank Kurt Braunlich for his advice on analysis tools. We thank the Love Lab for the helpful discussions on the project. We are grateful to members of Cognitive Brain Mapping Lab at RIKEN BSI for sharing natural images used in this study.

